# Using Haplotype and QTL Analysis to Fix Favorable Alleles in Diploid Potato Breeding

**DOI:** 10.1101/2022.11.09.515871

**Authors:** Lin Song, Jeffrey B. Endelman

**Affiliations:** Department of Horticulture, University of Wisconsin-Madison, Madison, WI 53706, USA

## Abstract

At present, the potato of international commerce is autotetraploid, and the complexity of this genetic system creates limitations for breeding. Diploid potato breeding has long been used for population improvement, and thanks to improved understanding of the genetics of gametophytic self-incompatibility, there is now sustained interest in the development of uniform F_1_ hybrid varieties based on inbred parents. We report here on the use of haplotype and QTL analysis in a modified backcrossing (BC) scheme, using primary dihaploids of *S*.*tuberosum* as the recurrent parental background. In Cycle 1 we selected XD3-36, a self-fertile F_2_ clone homozygous for the self-compatibility gene *Sli*. Signatures of gametic and zygotic selection were observed at multiple loci in the F_2_ generation, including *Sli*. In the BC_1_ cycle, an F_1_ population derived from XD3-36 showed a bimodal response for vine maturity, which led to the identification of late vs. early alleles in XD3-36 for the gene *StCDF1* (*Cycling DOF Factor 1*). Greenhouse phenotypes and haplotype analysis were used to select a vigorous and self-fertile F_2_ individual with 43% homozygosity, including for *Sli* and the early-maturing allele *StCDF1*.*3*. Partially inbred lines from the BC_1_ and BC_2_ cycles have been used to initiate new cycles of selection, with the goal of reaching higher homozygosity while maintaining plant vigor, fertility, and yield.

**Core Ideas:**

1. Partially inbred, diploid potato lines were developed for transitioning to an inbred-hybrid breeding system.
2. Multi-generational linkage analysis was used to track and fix favorable alleles without haplotype-specific markers.
3. Signatures of gametic and zygotic selection were detected by maximum likelihood.

## Introduction

In the 20th century, worldwide production and breeding of potato (*Solanum tuberosum* L.) was focused on autotetraploid (2n=4x=48) germplasm. During this time, there was also significant “pre-breeding” at the diploid level, primarily to facilitate the use of wild and cultivated germplasm from the Andean region of South America, where the potato was first domesticated. The culmination of diploid breeding was the transfer of beneficial alleles into tetraploid germplasm through 2x-4x crosses (i.e., unilateral sexual polyploidization), rather than clonal selection for variety release (Hougas & Peloquin, 1958; Chase, 1963) Inbreeding depression was well known in diploid potato (de Jong & Rowe, 1971), and tetraploidy offered more opportunities for complementation; this was called selection for “maximum heterozygosity” (Bingham, 1980). Based on this prevailing wisdom, 20^th^ century efforts to develop potato cultivars that can be propagated sexually (i.e., by “true” potato seed, TPS) utilized tetraploid rather than diploid germplasm (Golmirzaie et al., 1994).

*S. tuberosum* Group Andigenum diploids and many wild species exhibit gametic self-incompatibility (SI), in which S-RNase expressed in the pistil inhibits the growth of self-pollen tubes (Kubo et al., 2010). For nonself pollen, the S-RNase is targeted for degradation by F-box proteins, creating sexual compatibility. Despite the widespread presence of SI in diploid potato, self-compatible clones have been recognized and studied (Cipar, 1964; Olsder & Hermsen, 1976). Hosaka and Hanneman (1998) mapped the genetic locus underlying this self-compatibility, named *Sli* for S-locus inhibitor, to potato chromosome 12; by contrast, the potato S-locus is on chromosome 1. Map-based cloning has shown *Sli* encodes an F-box protein (Eggers et al., 2021; Ma et al., 2021).

Genetic understanding of self-compatibility has led to a paradigm shift in diploid potato breeding, commonly described as “Potato 2.0” (Stokstad, 2019). No longer limited to population improvement, diploids are being used to create inbred lines and F_1_ hybrid varieties that may eventually replace tetraploids (Phumichai et al., 2005; Lindhout et al., 2011). A diploid, inbred-hybrid breeding system offers many advantages to the current breeding system in potato: it takes less time to fix favorable alleles; marker-assisted backcrossing is possible; there is greater genetic variance for selection; and heterosis can be exploited systematically (Jansky et al., 2016). As mentioned already, inbreeding depression is a significant obstacle to realizing these goals, but compared to previous generations of breeders, genomics and computational tools are now available to expedite the identification and elimination of deleterious alleles (Zhang et al., 2019, 2021).

The University of Wisconsin-Madison potato breeding program was initiated in the 1930’s and has released a number of commercially successful tetraploid varieties during its history, particularly for the round white, potato chip market. Between 2016 and 2018, elite clones from the program were crossed as female parents with the haploid inducer IVP101 (Hutten et al., 1993) to generate dihaploid (diploid haploid, DH) founders for breeding. After screening hundreds of dihaploids under greenhouse conditions for vigor and female fertility, a handful have been used in a generalized backcrossing scheme (Fig. 1), to introduce *Sli* and other desirable traits into a more elite background. We report here on the outcomes of this breeding effort. A distinguishing feature of our approach has been the use of multi-generational linkage analysis to track identical-by-descent (IBD) haplotypes from the founders (Zheng et al., 2015). This allowed us to make rapid progress for fixation of *Sli* and early maturity at *CDF1* (*Cycling DOF Factor 1;* Kloosterman et al., 2013), even in the absence of haplotype-specific markers.

**Figure 1.**
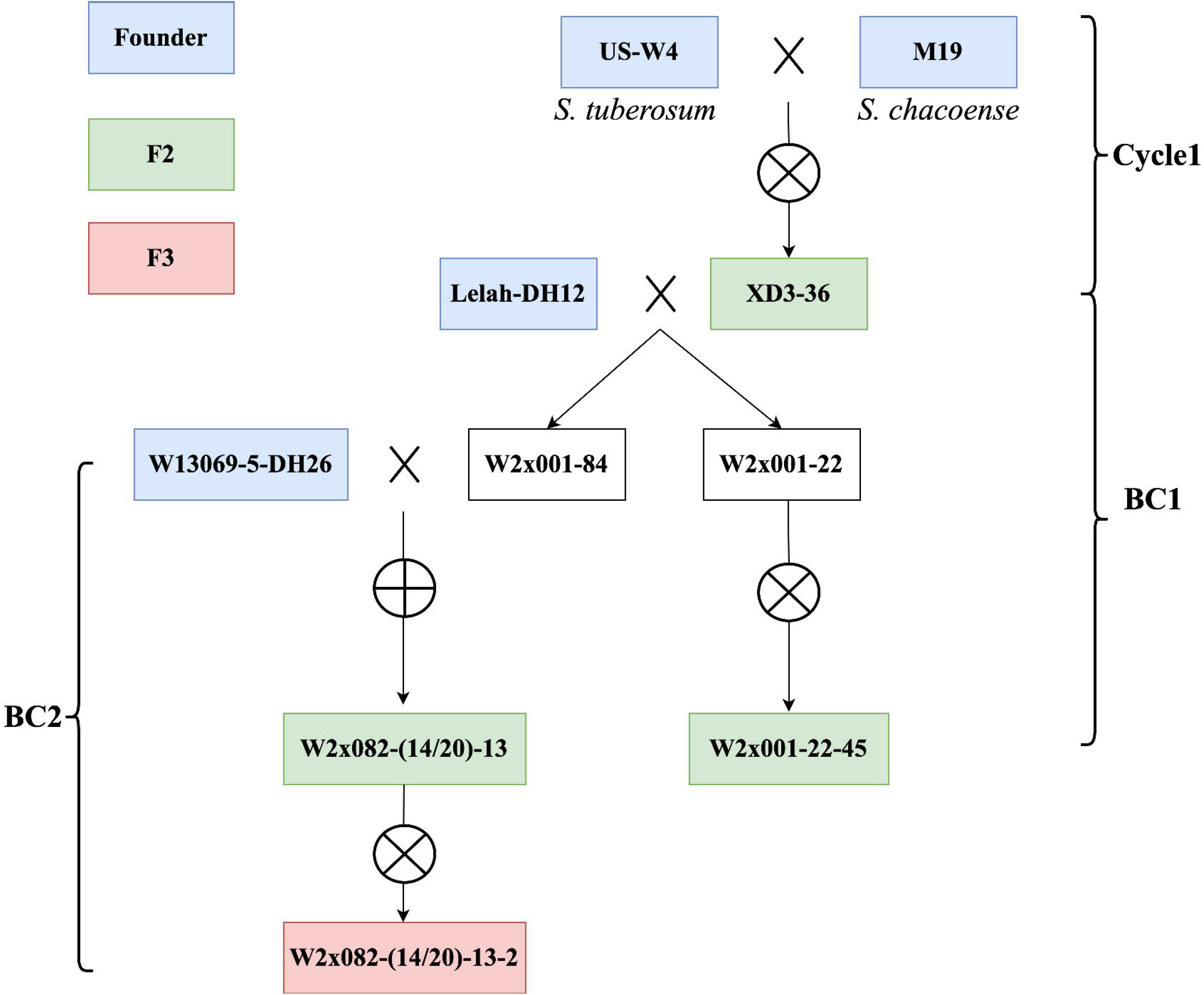
Breeding scheme to develop partially inbred lines fixed for favorable alleles at key loci. Standard nomenclature is used: [Cross]-[F1]-[F2]-[F3]. When sib-mating of two individuals was used instead of selfing, the naming convention was (ID1/ID2).

## Materials and Methods

### Nomenclature

Germplasm created during this research was named following the convention that a dash indicates generations separated by one meiosis during inbreeding, e.g., [Cross]-[F1]-[F2]-[F3]. Dihaploid progeny of tetraploid clones are labeled [Clone]-DH[ID]. The founder US-W4 (Peloquin & Hougas, 1960) does not follow this convention because it is legacy germplasm. The prefix “W2x” indicates “Wisconsin diploid” germplasm.

### Phenotyping

Unreplicated greenhouse experiments were conducted at the Walnut Street facility at the University of Wisconsin-Madison (Madison, WI). True potato seeds (TPS) were soaked for 24 h in 1500 ppm Gibberellic Acid to break dormancy before sowing in flats. Seedlings were transplanted into 3.8L pots approximately 28 days after planting (DAP). Environmental conditions were a 16h day/8h night photoperiod, with daytime temperatures 18–22°C and nighttime temperatures 16–20°C. Five traits were measured: pollen shed, vine maturity, stolon production, tuber yield, and seed (TPS) yield. Pollen shed was scored as a binary trait based on visual observation after self-pollinating at least 10 flowers per plant. Vine maturity was visually rated on a scale of 1 (early) to 9 (late) at 144 DAP. Stolon production was visually rated on a scale of 1 (few) to 5 (abundant) at harvest 150 DAP. Tuber yield was the total tuber weight (g) per plant. Seed yield was the number of seeds per plant.

Field evaluation of the BC_1_F_1_ population occurred at the UW-Madison Hancock Agricultural Research Station (HARS) (Hancock, WI) in 2019. A partially replicated, incomplete block design was used, with a single plot for 89 progeny and two plots for 9 progeny and both parents. Eight seed pieces per plot were planted April 30 and harvested 122 DAP, two weeks after vine dessication with diquat. Fertilization, irrigation, and pest management followed standard practice (Bussan et al., 2015). Vine maturity was visually rated using the same 1–9 scale at 100 DAP. Yield was calculated on a per plant basis by dividing the plot weight by the stand count. Size A tubers (diameter > 4.8 cm) were separated using a chain grader to report the A-size proportion on a weight basis.

Ten tubers were stored at 7°C, 95% RH for 3 mo before measuring post-harvest quality traits. Specific gravity was measured based on underwater weight (Wang et al., 2017). Fry color was measured on 1 mm chip slices, fried for 2 min and 10 s at 360F. Chips were crushed before measuring reflectance on the Hunter Lightness scale (L) with a HunterLab D25NC colorimeter (Reston, VA).

Five tubers were stored at 12°C, 95% RH for 10 weeks before the start of a 16-week experiment to measure tuber dormancy. Every 2 weeks, tubers were individually scored using a 3-point scale for the length of sprouts: 0 = none, 0.5 = less than 2mm, and 1 = above 2mm. The average of the five tubers was the dormancy score for each plot, and the relative area under the sprout vs. time curve (AUC) was calculated on a 0-1 scale.

### Genotyping

Two different platforms were used to obtain genome-wide markers in this project. For tracking IBD haplotypes across the breeding cycles, we used version 3 of the potato SNP array (Felcher et al., 2012; Vos et al., 2015), which generated 10,322 markers. As part of an experimental project on genotyping-by-sequencing (GBS) in potato, the BC_1_F_1_ population was genotyped at the University of Minnesota Genomics Center (UMGC) with a two-enzyme (MspI + PstI) protocol (Poland et al., 2012). Approximately 130M reads were obtained with the Illumina NextSeq 1×150 bp platform, and variant discovery was performed by the genotyping service provider using FreeBayes (Garrison & Marth, 2012) and the DMv4.03 reference genome (Potato Genome Sequencing Consortium, 2011; Sharma et al., 2013). Variant filtering was performed using custom R scripts (R Core Team, 2022). Only bi-allelic SNPs with a minimum sample depth of 10 reads, less than10% missing data, and minor allele frequency > 0.05 were retained, yielding 7673 markers. Genotype calls were made using R package *updog* with the “f1” model to account for allelic bias and overdispersion (Gerard et al., 2018).

KASP genotyping with marker *Sli_898* (Clot et al., 2020; Kaiser et al., 2021) was used to confirm *Sli* genotypes inferred from the haplotype analysis. The protocol of Kaiser et al. (2021) was followed using the KASP v4.0 2x standard ROX Master Mix (LGC Genomics, Beverly, MA) and detection with the Bio-Rad CFX96 equipment.

Whole-genome sequencing of the BC_1_F_2_ individual W2×001-22-45 utilized the NovaSeq 2×150 flow cell (University of Minnesota), with a yield of 376M paired reads. Reads were aligned with BWA-MEM (Li, 2013) to the *CDF1*.*1_scaffold1389* and *CDF1*.*3_scaffold390* alleles from the Atlantic reference genome (Hoopes et al., 2022) and then filtered to remove alignments with fewer than 10 bp on both sides of the transposon insertion site (Caraza-Harter & Endelman, 2022). Only alignments to the *CDF1*.*3* reference were detected, confirming homozygosity for this allele.

### Genetic analysis

Multi-generational tracking of IBD haplotypes was conducted using the SNP array marker data and the software RABBIT (Zheng et al. 2015; 2018), with marker order based on the DMv6.1 reference genome (Pham et al., 2020). First, we analyzed the Cycle 1 genotypes, using the RABBIT MagicImpute function to phase US-W4 and M19 as outbred founders. This analysis generated a phased genotype for their F_1_ offspring XD3, which was then used as one of three founders—the others being Lelah-DH12 and W13069-DH26 (Fig. 1)—to analyze all three breeding cycles together (Files S2 and S3). Based on the haplotype reconstruction of XD3 in terms of M19 and US-W4, all individuals were reconstructed in terms of 6 founder haplotypes: M19, US-W4, Lelah-DH12.1, Lelah-DH12.2, W13069-DH26.1, W13069-DH26.1, where “.1” and “.2” refer to the two haplotypes in an outbred diploid. The maximum posterior genotypes are in File S4.

Signatures of gametic and zygotic selection in the Cycle 1 F_2_ population were detected using maximum likelihood (ML). Two models of gametic selection were considered, based on whether one (gametic1) or both (gametic2) sexes experience selection. The selection coefficient *s* quantifies the strength of selection, with positive (negative) values representing selection against the A (B) allele. Two models of zygotic selection were considered, based on whether one or both homozygotes experience selection. Under the zygotic1 model, positive (negative) values of *s* represent selection against AA (BB). Under the zygotic2 model, positive (negative) values of *s* represent selection against (for) the homozygotes. By specifying that *s* equals the sum of the absolute deviations between the expected (without selection) and observed frequencies, we derived the expected frequency *p* of the three possible genotypes (AA, AB, BB) for the four selection models (Table 1). If *N*_*AA*_, *N*_*AB*_, *N*_*BB*_ represent the observed counts of each genotype, the log-likelihood (LL) of this outcome is *N*_*AA*_ log *p*_*AA*_ + *N*_*AB*_ log *p*_*AB*_ + *N*_*BB*_ log *p*_*BB*_. R function *optimize* was used to identify the ML solution for *s* for each model, and the model with the highest LL was selected for each marker (File S5). The likelihood ratio (i.e., Wilks) test was used to compute the p-value for the null hypothesis of no selection: *s* = 0. A Bonferroni-corrected significance threshold of 0.05/*m* was used for detection, where *m* is the total number of markers.

**Table 1.**
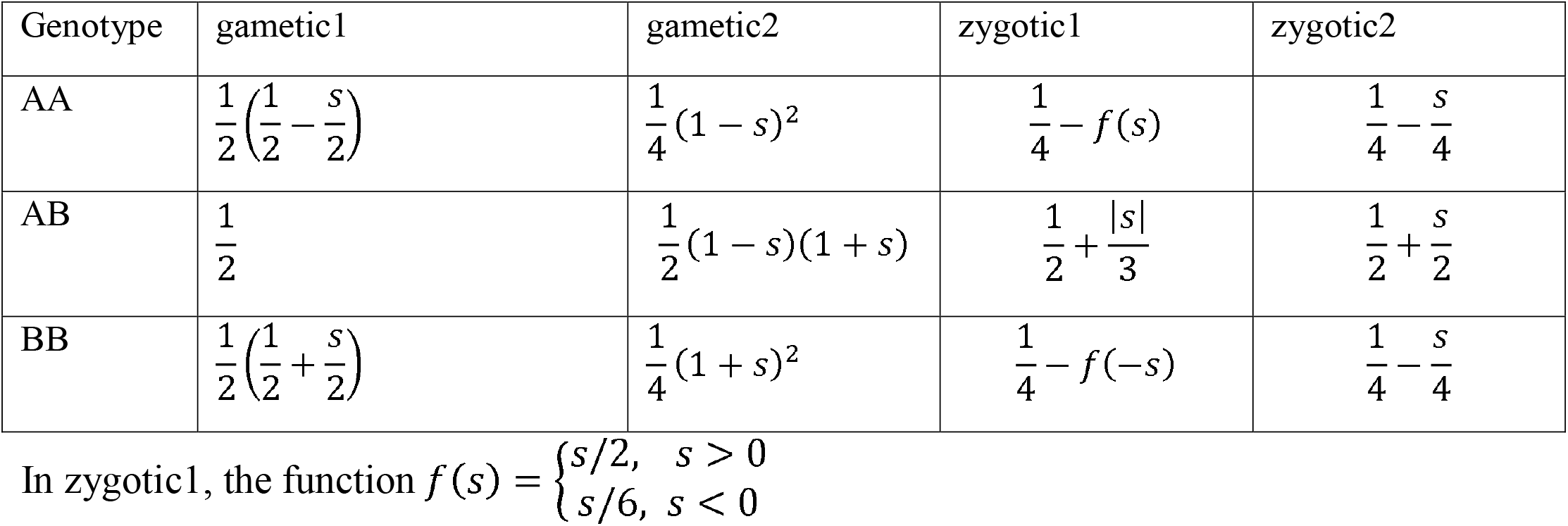
Genotype frequencies under gametic and zygotic models of selection in F_2_ populations.

QTL mapping was conducted for the BC_1_F_1_ W2×001 population with R package diaQTL (Amadeu et al., 2021). The recommended settings from the package tutorial were used for the number of iterations, and the discovery threshold for the single QTL scan was based on a genome-wide significance level _α_ = 0.05. Phasing of the outbred parents and haplotype reconstruction of 132 progeny were performed using *PolyOrigin* (Zheng et al., 2021; File S6 contains the input marker data), with the following parameters: isphysmap=true, recomrate=1.25, refineorder=false, refinemap=true. PolyOrigin did not produce sensible results for chromosome 11 because it was completely homozygous in one parent, so RABBIT was used instead. The genotype probability input file for diaQTL (File S7) was generated from the PolyOrigin and RABBIT outputs using the functions *convert_polyorigin* and *convert_rabbit*, respectively, in the diaQTL package.

The phenotypes for QTL mapping (File S8) come from the unreplicated greenhouse and partially replicated field experiments described above. For traits in the field trial, fixed effect estimates for genotype (*g*_*i*_) were used as the response variable, based on Eq. 1:

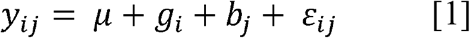

In Eq.1, *y*_*ij*_ is the response for genotype *i* in block *j, μ* is the population mean, *b*_*j*_ is the fixed effect for block, and *ε*_*ij*_ is the residual with variance σ_ε_^2^. Variance components were estimated using ASReml-R (Butler et al., 2018). Eq. 1 was also used to estimate broad-sense heritability (H^2^) on a plot basis by treating the genotype effect as random with variance σ_g_^2^:

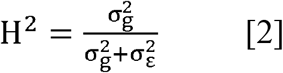

## Results

### Cycle 1

Cycle 1 was initiated with the goal of identifying a fertile F_2_ clone homozygous for *Sli*, to be used as the male parent in a generalized backcrossing scheme with *S. tuberosum* dihaploids (Fig. 1). The grandparents of the F_2_ population were an “heirloom” *S. tuberosum* dihaploid US-W4 (Peloquin & Hougas, 1960) and an inbred clone M19 from the wild species *S. chacoense* (Fulladolsa et al., 2019). At that time (2018), it was believed that introgression of *Sli* from *S. chacoense* into *S. tuberosum* was necessary, and thus our strategy was to identify self-fertile F_2_ clones homozygous for the M19 haplotype in the vicinity of the published location of *Sli* on chromosome 12. To our surprise, there were no offspring homozygous for M19 in this region (Fig. 2). The ratio of US-W4 homozygotes to heterozygotes was approximately 1:1, which is consistent with self-fertilization only by pollen containing *Sli* on the US-W4 haplotype. This interpretation was corroborated by Clot et al. (2020), who identified kmers linked to *Sli* and reported their presence in US-W4, along with many other *S. tuberosum* clones.

**Figure 2.**
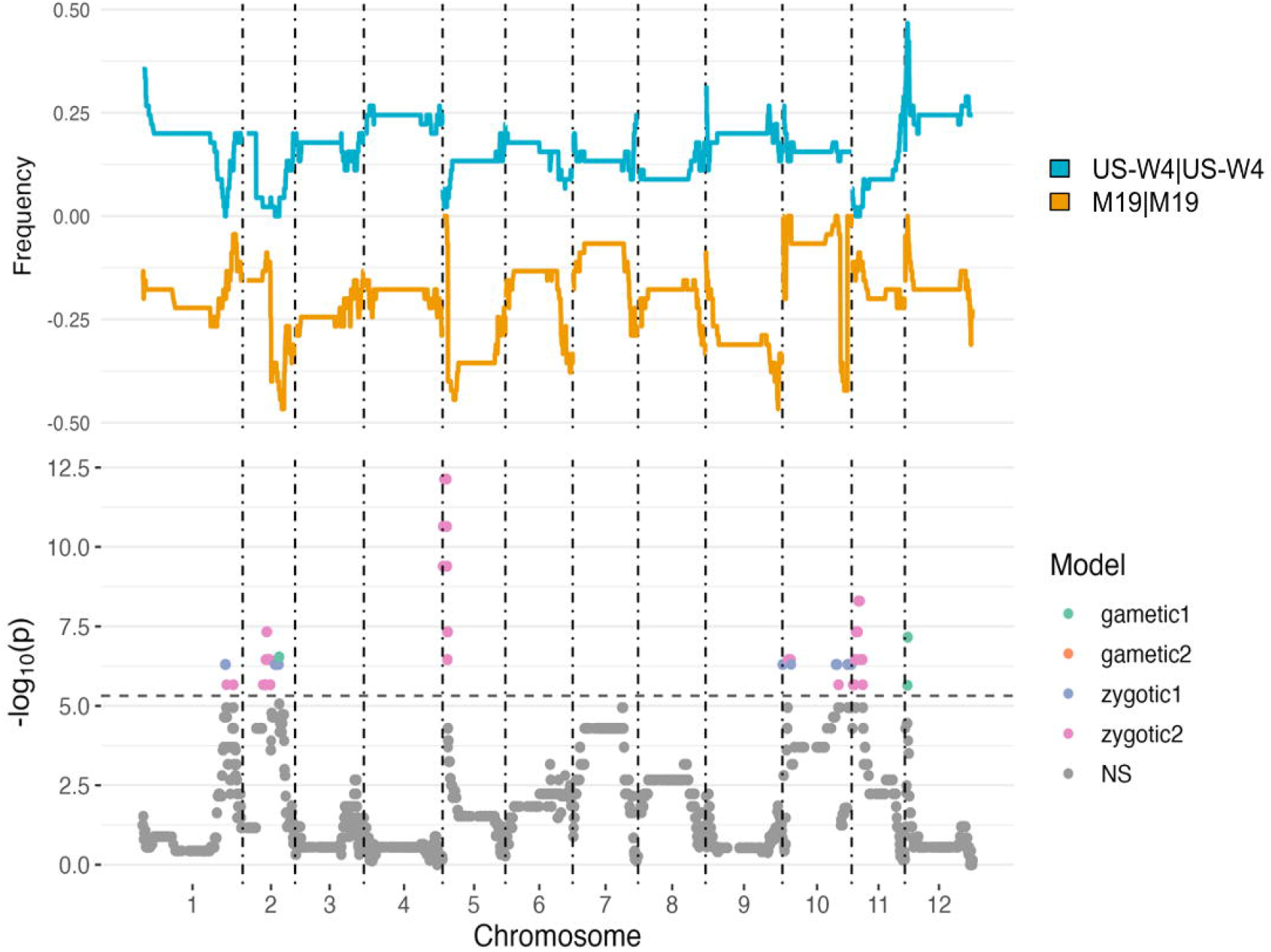
Signatures of selection in the F_2_ generation of Cycle 1. (Top) Homozygote frequencies. (Bottom) Hypothesis testing for zygotic and gametic selection at significance level α = 0.05, with Bonferroni correction for multiple testing.

Several other genomic regions displayed signatures of selection in the Cycle 1 F_2_ population (Fig. 2). Maximum likelihood was used to categorize distorted segregation into one of four possible selection models: gametic selection on one sex (gametic1), gametic selection on both sexes (gametic2), zygotic selection on one homozygote (zygotic1), and zygotic selection on both homozygotes (zygotic2). Zygotic selection against both homozygotes was the most common inference, although in some regions multiple selection models were significant (Table S1).

The F_2_ individual, XD3-36, was selected as a male parent for the first backcross (BC_1_) cycle based on its desirable combination of traits: good tuber yield (not recorded), seed production (340 seeds), and homozygosity for *Sli*. The *Sli* genotype was originally inferred based on haplotype analysis (Fig. 3) and later confirmed using KASP markers developed by Clot et al. (2020).

**Figure 3.**
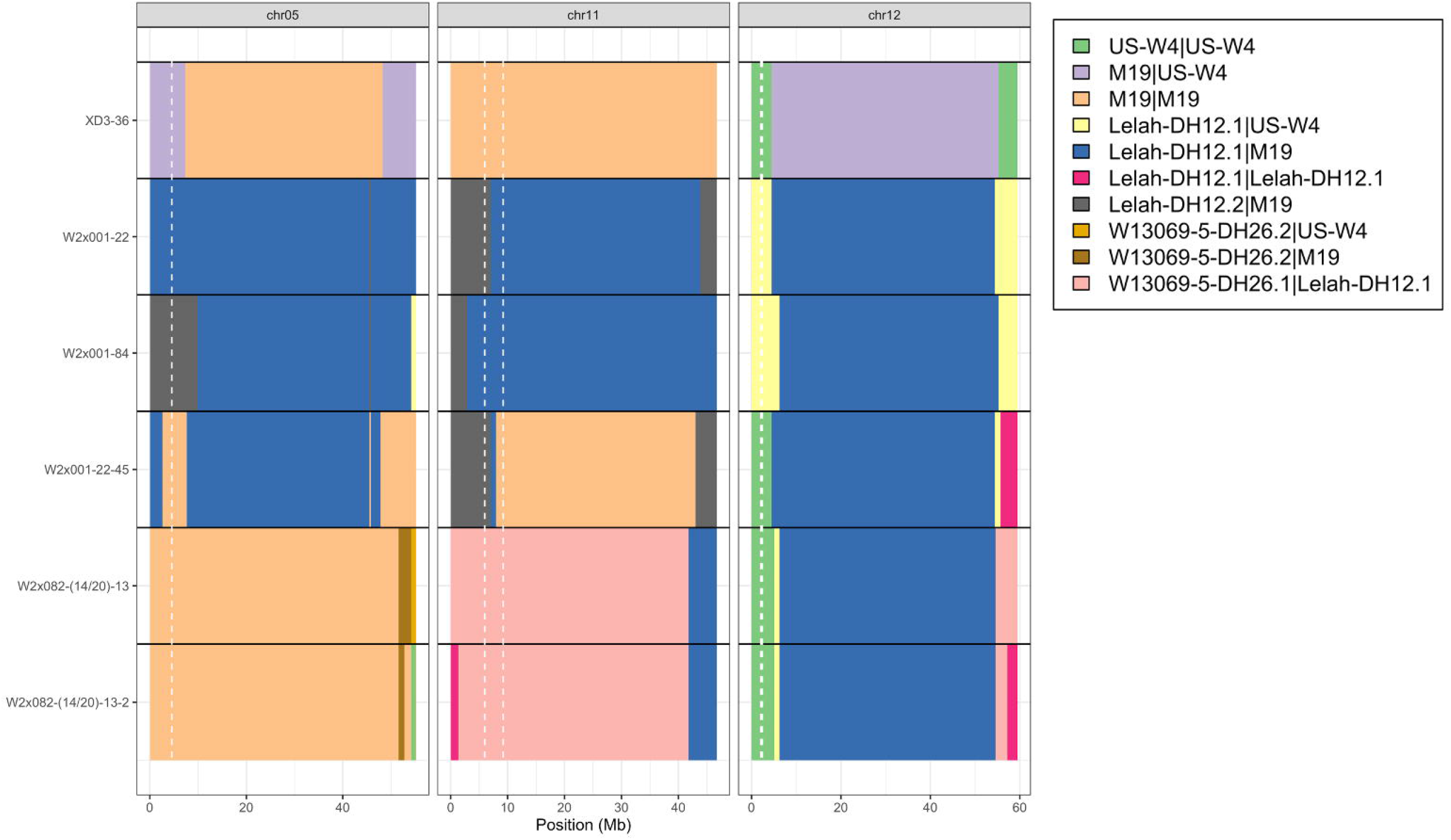
Haplotype reconstruction of key genotypes for chromosome 5, 11 and 12. White dashed lines represent the location of *CDF1* on chromosome 5, the fertility QTL on chromosome 11, and *Sli* on chromosome 12. All 12 chromosomes shown in Figure S4.

### Cycle BC_1_

Several BC_1_F_1_ populations were created from different dihaploid mothers, but population W2×001 from Lelah-DH12 was singled out for more intensive study. Visual ratings for vine maturity and stolon production were correlated and exhibited bimodal distributions (Fig. S1). There were 29 plants with abundant pollen shed, and 16 produced fruit upon selfing. The number of seeds among the self-fertile plants was skewed, with a range of 15 to 662 and median 119 (Fig. S1). Tuber yield ranged from 0 to 973g per plant, with a median of 370g. Tuber yield was significantly higher, by 245g (p = 6x 10-9), for the plants with pollen shed.

There were enough greenhouse tubers for 98 F_1_ progeny to conduct a partially replicated, clonal field trial in 2019. A number of agronomic and quality traits were measured, with broad-sense heritability on a plot basis between 0.56 (total yield) and 0.86 (tuber dormancy; Table 2). Unlike the greenhouse study, the distribution for vine maturity was not bimodal. Total yield per plant ranged from 0.19 to 1.41 kg (median 0.73), compared to 0.47 and 0.27 kg for the parents Lelah-DH12 and XD3-36, respectively. Specific gravity and fry color lightness, which are important traits for the potato chip market, were measured after 3 mo of storage. Spec. gravity ranged from 1.050 to 1.114 (median 1.082), and fry color ranged from 33.2 to 59.3 (median 48.9). Higher values of spec. gravity and fry color were positively correlated with each other (*r* = 0.59) and negatively correlated with tuber size (*r* = -0.52 and -0.42, respectively; Fig. S2).

**Table 2.** Broad-sense heritability estimates (plot basis) from the field trial of the W2×001 BC_1_F_1_ population.

| Trait | H<sup>2</sup> |
| --- | --- |
| Vine Maturity | 0.65 |
| Tuber Yield | 0.56 |
| Tuber Size (% A) | 0.67 |
| Tuber Appearance | 0.66 |
| Specific Gravity | 0.74 |
| Fry Color | 0.61 |
| Tuber Dormancy | 0.86 |

Genotyping-by-sequencing of the F_1_ population for QTL analysis led to creation of a genetic map with 7497 markers and 1553 cM. For vine maturity, stolon production, and tuber dormancy, a QTL in the vicinity of *CDF1* on potato chromosome 5 was detected, which explained 53, 34, and 23% of the variance for those three traits, respectively (Table 3). The estimated parental haplotype effects indicated the *CDF1* allele inherited from M19 was significantly earlier than the allele from US-W4 (Table 3, Fig. 4). There was no significant difference between the two haplotypes from Lelah-DH12 at *CDF1*, which suggests they carry the same allele. A binary trait locus, or BTL, was detected for pollen shed on chromosome 11, explaining 43% of the variance. The parental haplotype effects indicate this BTL is the result of allelic differences in Lelah-DH12, with the favorable allele for fertility on haplotype Lelah-DH12.1. Additional QTL for vine maturity on chromosome 1 and tuber dormancy on chromosome 7 explained 15-19%.

**Table 3.**
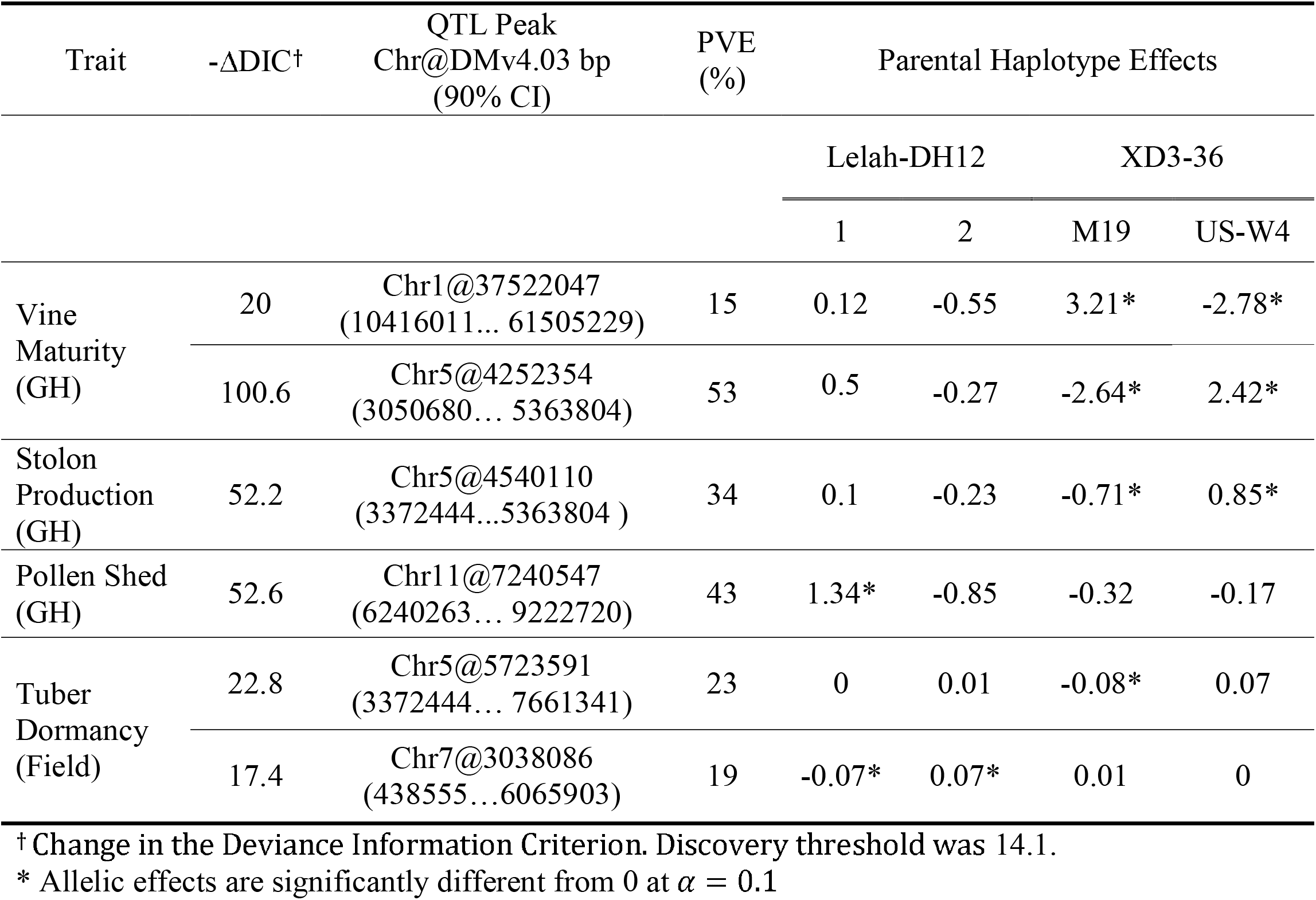
Quantitative trait loci (QTL) for the W2×001 BC_1_F_1_ population.

**Figure 4.**
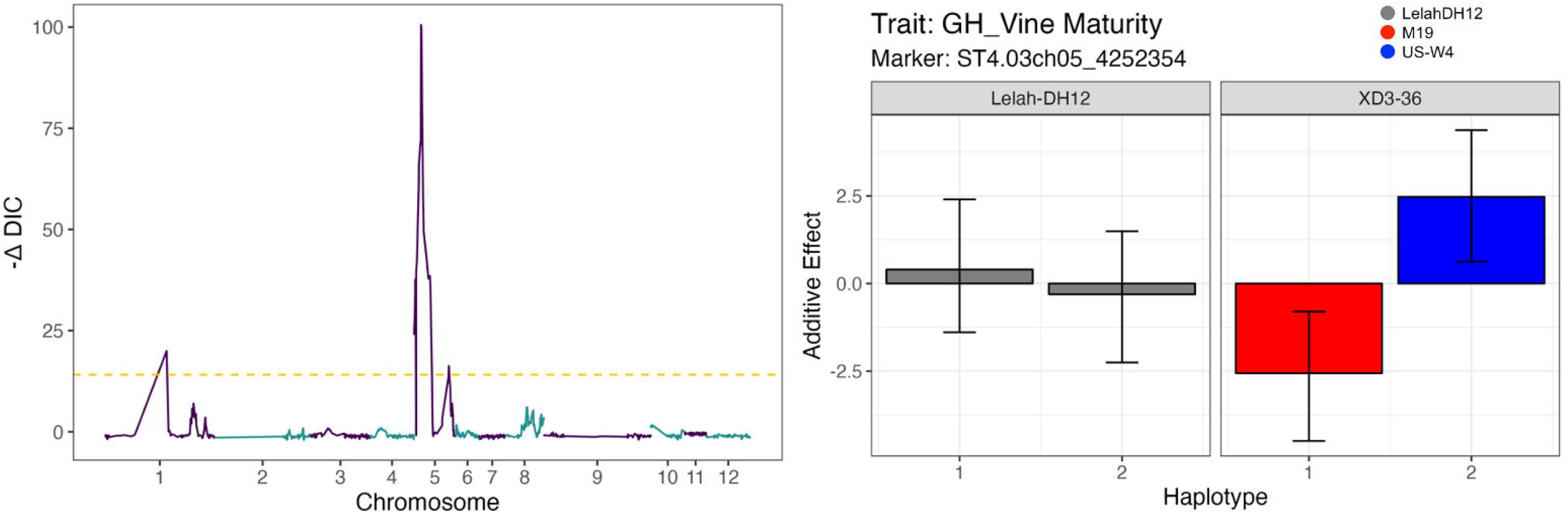
Genetic mapping of greenhouse vine maturity in the BC_1_F_1_ population. (Left) Single QTL genome scan. (Right) Parental haplotype effects for the QTL on chromosome 5. Higher trait values represent later maturity.

Based on the number of selfed seeds and tuber yields of the F_1_ progeny, 17 F_2_ families from W2×001 were selected for greenhouse evaluation. One of the best families, in terms of plant vigor and female self-fertility, was derived from W2×001-22. Haplotype analysis of the F_2_ population revealed no homozygotes of the Lelah-DH12 haplotype at the *Sli* locus (Fig. S3), which indicates this haplotype in W2×001-22 did not carry *Sli*. To achieve homozygosity for early maturity and *Sli*, we selected F_2_ progeny homozygous for the US-W4 haplotype at *Sli* and homozygous for the M19 haplotype at *CDF1*. One particular F_2_ individual, W2×001-22-45, met these criteria and had good tuber yield and self-fertility (Fig. 3 and 5). Homozygosity varied by chromosome from a low of 2% on chr01 to 99.9% on chr04, with a genome-wide average of 43% (Table 4 and Fig. S4).

**Table 4.** Homozygosity percentages (based on DMv6.1 bp) for partially inbred clones from the modified backcrossing scheme.

| Chr | XD3-36<br>Cycle 1 | W2x001-22-45<br>Cycle BC<sub>1</sub> | W2x082-(14/20)-13<br>Cycle BC<sub>2</sub> | W2x082-(14/20)-13-2<br>Cycle BC<sub>2</sub> |
| --- | --- | --- | --- | --- |
| 1 | 75.1 | 2.2 | 0.0 | 0.0 |
| 2 | 72.5 | 63.7 | 0.9 | 36.4 |
| 3 | 0.0 | 50.1 | 6.5 | 72.6 |
| 4 | 73.5 | 99.9 | 2.5 | 88.8 |
| 5 | 73.8 | 22.1 | 93.3 | 96.5 |
| 6 | 17.0 | 22.1 | 0.0 | 99.2 |
| 7 | 4.2 | 7.6 | 0.0 | 0.0 |
| 8 | 7.1 | 71.6 | 0.0 | 9.2 |
| 9 | 3.4 | 16.3 | 6.3 | 12.8 |
| 10 | 7.0 | 74.2 | 10.8 | 92.7 |
| 11 | 100.0 | 75.5 | 0.0 | 1.8 |
| 12 | 13.8 | 13.4 | 8.1 | 11.9 |
| Average | 37.3 | 43.2 | 10.7 | 43.5 |

**Figure 5.**
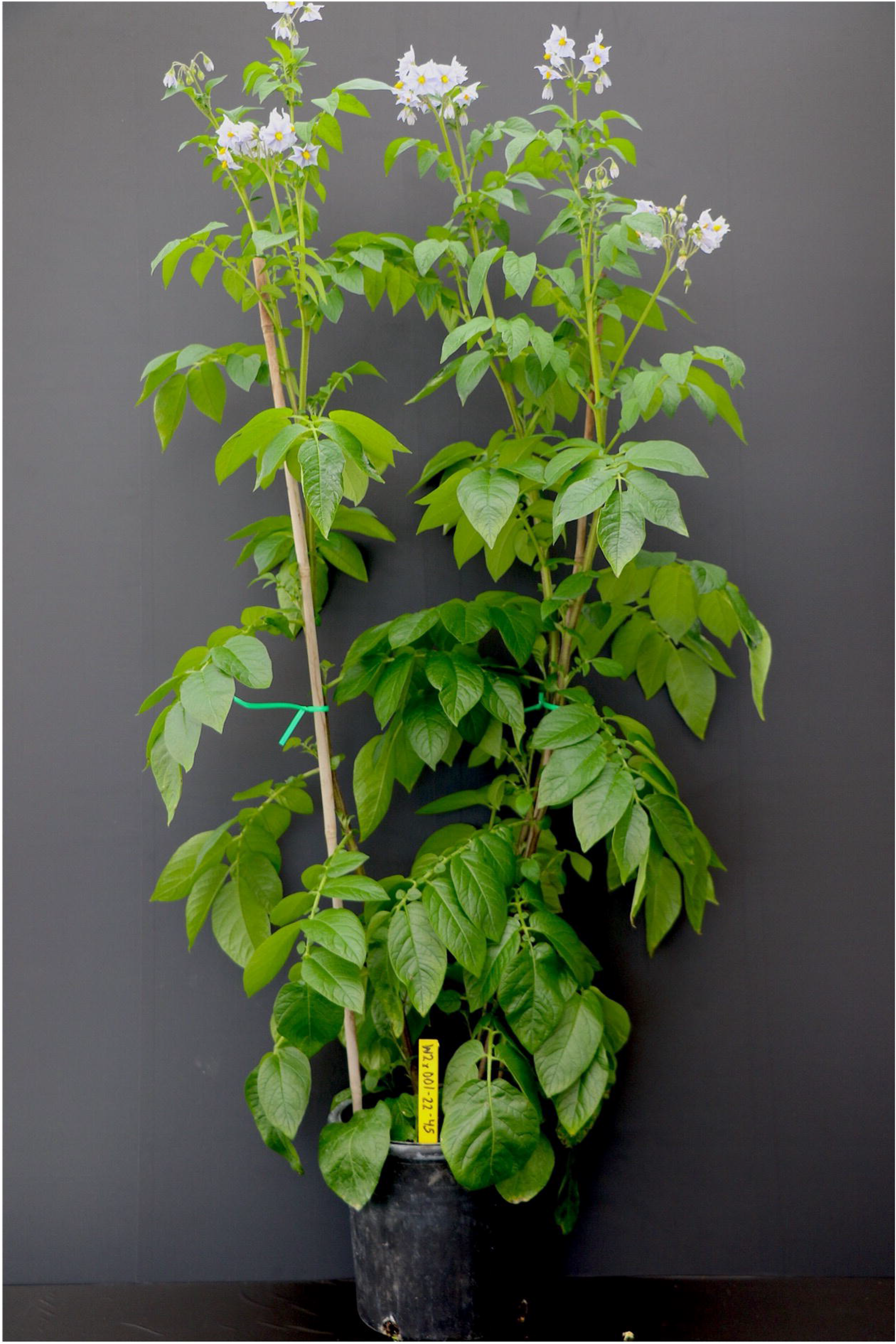
The BC_1_F_2_ individual W2×001-22-45, which was selected based on vigor, self-fertility, and tuber yield in greenhouse experiments. It is homozygous for *Sli* and *CDF1*.*3*.

Whole-genome sequencing of W2×001-22-45 was used to determine which *CDF1* allele is present. As expected, only one allele was detected: *CDF1*.*3*, which is the earliest known allele and encodes a truncated protein without the FKF1 binding domain (Kloosterman et al., 2013).

### Cycle BC_2_

Cycle BC_2_ was initiated by using two superior BC_1_F_1_ individuals, W2×001-22 and W2×001-84, as pollen donors to fertilize *S. tuberosum* dihaploids. Both selfing and sib-mating of BC_2_F_1_ individuals were used for inbreeding, and one F_2_ population derived from sib-mating W2×082-14 and W2×082-20 had particularly good characteristics (Fig. 1). Haplotype analysis in the F_2_ generation enabled genetic selection for homozygosity at *CDF1* and *Sli* (Fig. 3) and phenotypic selection for tuber and (selfed) F_3_ seed yield. The top F_3_ population derived from the F_2_ individual W2×082-(14/20)-13, which was later determined to be only 11% homozygous—well below the expected value of 25% for a sib-mated F_2_. Tuber yield and homozygosity were inversely related (*r* = -0.40, *p* < 0.05) in the F_3_ population (Fig. S5), while seed and tuber yields were positively correlated (*r* = 0.55, *p* < 0.05). Our top F_3_ selection, W2×082-(14/20)-13-2 (Fig. S6), had a bimodal distribution for homozygosity across the 12 chromosomes, with 4 chromosomes above 85% and 6 below 15% (Table 4). The genome-wide average of 43% homozygosity for W2×082-(14/20)-13-2 was comparable to the top selection in the BC_1_F_2_ generation, W2×001-22-45.

## Discussion

A potential challenge with the development of inbred-hybrid varieties in potato is competition between the sexual and asexual reproductive organs as sinks for assimilates (Almekinders & Struik, 1996). The potato tuberization pathway is the result of neofunctionalization of the flowering pathway, with the phloem-mobile signal for tuberization SP6A homologous to the florigen signal SP3D (Navarro et al., 2011; Abelenda et al., 2014). Both proteins are regulated by CDF1, and the two processes typically occur contemporaneously. To promote flowering, it is common practice in potato breeding to plant mother tubers on a brick for crossing, so that the soil can be washed away after the roots are established and daughter tubers removed (Thijn, 1954). However, not all genotypes respond to this treatment (Plantenga et al., 2019), and we observed a *positive* correlation between seed and tuber yield in both the BC_1_F_1_ and BC_2_F_3_ generations (Figures S2 and S5). Since both traits are likely to benefit from increased plant vigor, this may be expected when there is inbreeding depression. More research is needed to understand the conditions under which flowering and tuberization are antagonistic.

One of our goals during inbreeding was to select for homozygosity of the haplotype containing an early maturing allele at *CDF1*. It was only years later that we determined the allele was *CDF1*.*3* from whole-genome sequencing. Ramírez Gonzales et al. (2021) also generated diploids homozygous for *CDF1*.*3*, reporting they were “extremely weak with a stunted growth habit.” This observation was rationalized based on their discovery that the transposon insertion in *CDF1*.*3* disrupts the long non-coding RNA *StFLORE*, which is anti-sense to the *CDF1* transcript and helps to regulate stomatal opening. We did not observe a deleterious phenotype associated with homozygosity of *CDF1*.*3* in either the BC_1_ or BC_2_ cycle, so further research is needed to understand whether this is due to compensatory alleles in our germplasm. The W2×001-22-45 clone has been deposited with the US Potato Genebank (accession id ‘BS 451’) for other breeders and researchers to use.

Historically, the conventional wisdom in potato breeding was that diploid germplasm was useful for population improvement but not for the release of commercial varieties, primarily because of limitations for tuber size and yield. Following a decade of breeding for inbred diploids, a yield gap between diploid F_1_ hybrids and tetraploids was still evident in the publication by Stockem et al. (2020). This result is not too surprising given the significant inbreeding depression observed in potato (de Jong & Rowe, 1971; Zhang et al., 2019). In the case of maize, it took multiple decades before commercially viable F_1_ hybrids were developed (Duvick, 2005), and a higher density of deleterious alleles is expected for cultivated potato compared to maize (Hardigan et al., 2017; Hoopes et al., 2022). On the bright side, the current study and Zhang et al. (2021) have shown that genetics and genomics can be used to guide and accelerate inbreeding. We remain optimistic that inbred-hybrid varieties will eventually replace tetraploid clones.

## Supplemental Material

File S1. Supplemental Table and Figures.

File S2. Marker genotype data for RABBIT.

File S3. Pedigree data for RABBIT.

File S4. RABBIT haplotype reconstruction results.

File S5. R function to detect signatures of selection in F_2_ populations.

File S6. Marker genotype data for W2×001 for PolyOrigin.

File S7. Genotype probabilities for W2×001 for diaQTL.

File S8. Phenotype data for W2×001 for diaQTL.

## Author Contributions

LS: Investigation, Formal analysis, Writing – original draft, review and editing. JBE: Supervision, Investigation, Formal analysis, Writing – review and editing.

## Acknowledgments

We thank Grace Christensen for assistance with tissue culture and other lab protocols, Peyton Sorensen for dihaploid extraction, staff from the Rhinelander and Hancock Agricultural Research Stations, the UW-Madison Walnut Street Greenhouse Facility, and the Douches Lab for *Sli* KASP reagents. Financial support was provided by the USDA National Institute of Food Agriculture Award 2019-51181-30021.

## Data Availability

SNP marker and phenotype data are provided as supplemental files. Whole-genome sequence data for W2×001-22-45 is available from the NCBI Sequence Read Archive under BioProject ID PRJNA898285 (https://www.ncbi.nlm.nih.gov/bioproject/898285).

## Abbreviations

BC: backcross
*CDF1*: *Cycling DOF Factor 1*
DH: dihaploid
*Sli*: *S-locus* inhibitor
SI: self-incompatibillity
TPS: true potato seed

