## Supplementary material for "Using Haplotype and QTL Analysis to Fix Favorable Alleles in Diploid Potato Breeding": File S1

**Table S1.** Genomic regions under selection in the Cycle 1 $F_2$ population.

| Chr | DMv6.1 bp | Gametic Selection | | Zygotic Selection | |
|---|---|---|---|---|---|
| | | one sex | both sex | one homozygote | both homozygotes |
| 1 | 73076873…73814220 | | | US-W4\|US-W4 | |
| | 74311673…74404151 | | | | ✓ |
| | 80280351…80411707 | | | | ✓ |
| 2 | 17740253…24423913 | | | | ✓ |
| | 28393167…31826370 | | | US-W4\|US-W4 | |
| | 31939524…32116945 | US-W4 | | | |
| 5 | 170730...4650207 | | | | ✓ |
| 10 | 94681…655677 | | | M19\|M19 | |
| | 4857510…7325042 | | | | ✓ |
| | 7615227 | | | M19\|M19 | |
| | 46831995…48176087 | | | M19\|M19 | |
| | 49098058…49469871 | | | | ✓ |
| | 57080825…573330314 | | | M19\|M19 | |
| | 60460277…60527066 | | | M19\|M19 | |
| 11 | 1062832...9702537 | | | | ✓ |
| 12 | 2189494…2429036 | M19 | | | |

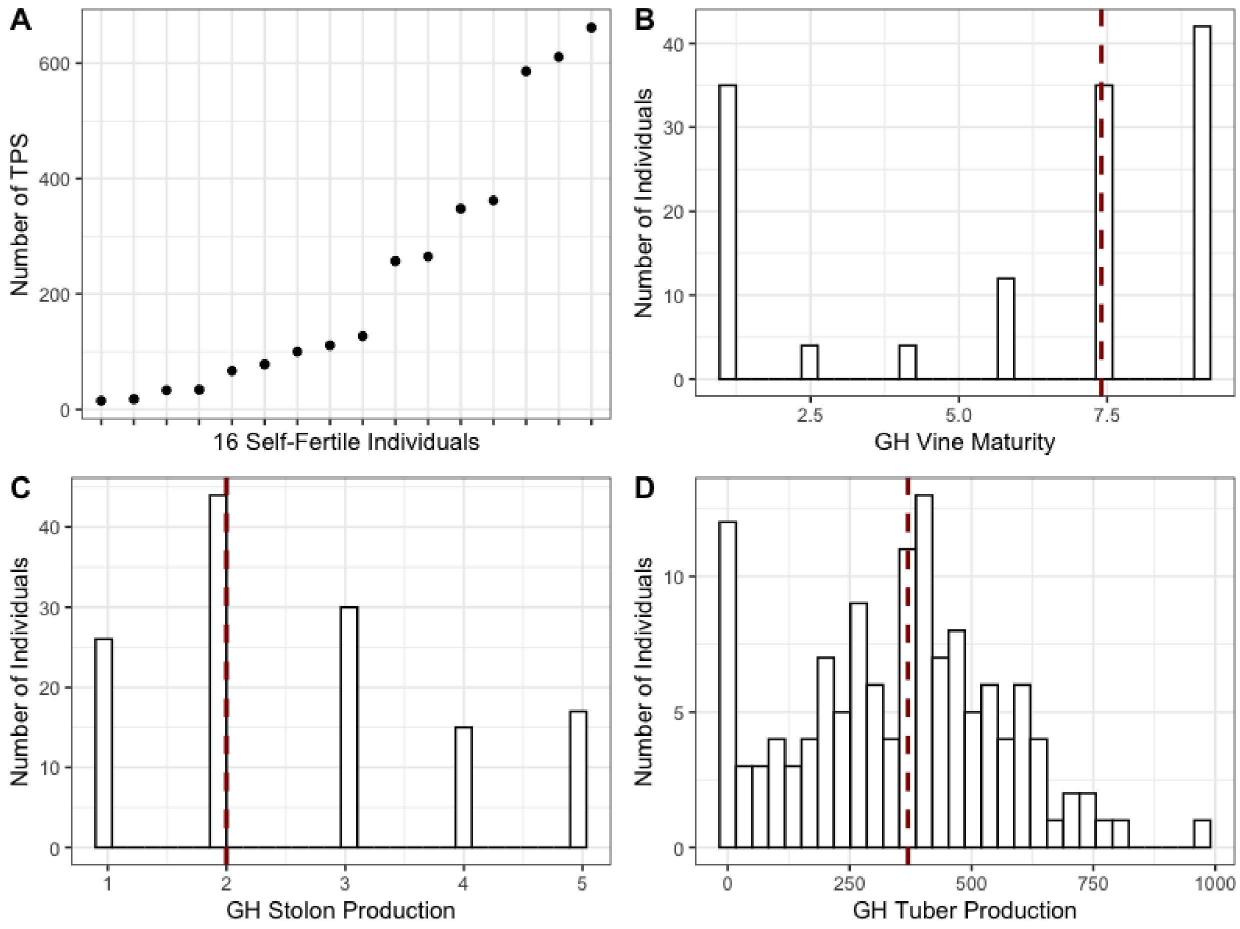


**Figure S1.** Greenhouse phenotypes for $BC_1F_1$ population: Number of true potato seeds per plant (A), GH Vine Maturity (B), GH Stolon Production (C), and GH Tuber Production (D). The Red dashed line represents the median of the trait.

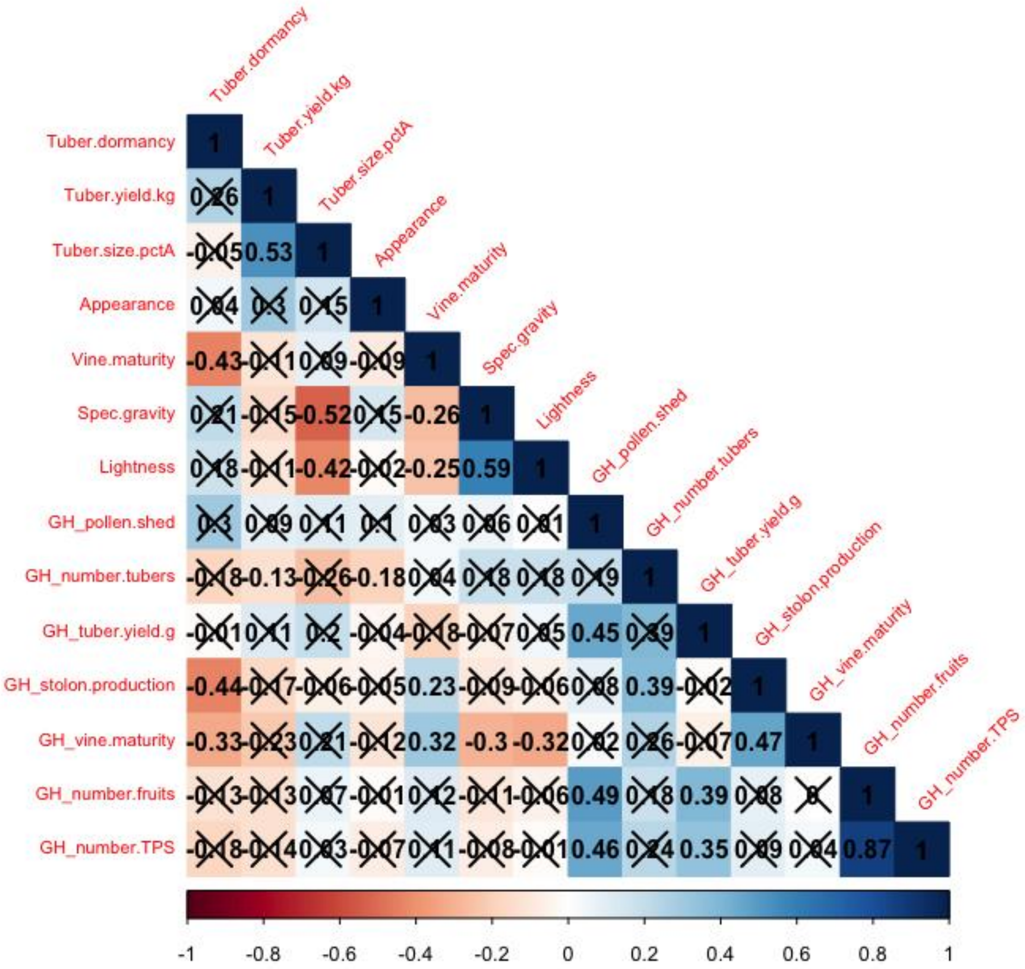


**Figure S2.** Correlation of traits evaluated in the greenhouse (GH prefix) and field for the $BC_1F_1$ population. The correlation coefficients are calculated using Pearson's R for continuous-continuous cases, correlation ratio for categorical-continuous cases, and Cramer's V for categorical-categorical cases. "X" denotes a non-significant correlation at $\alpha$=0.05.

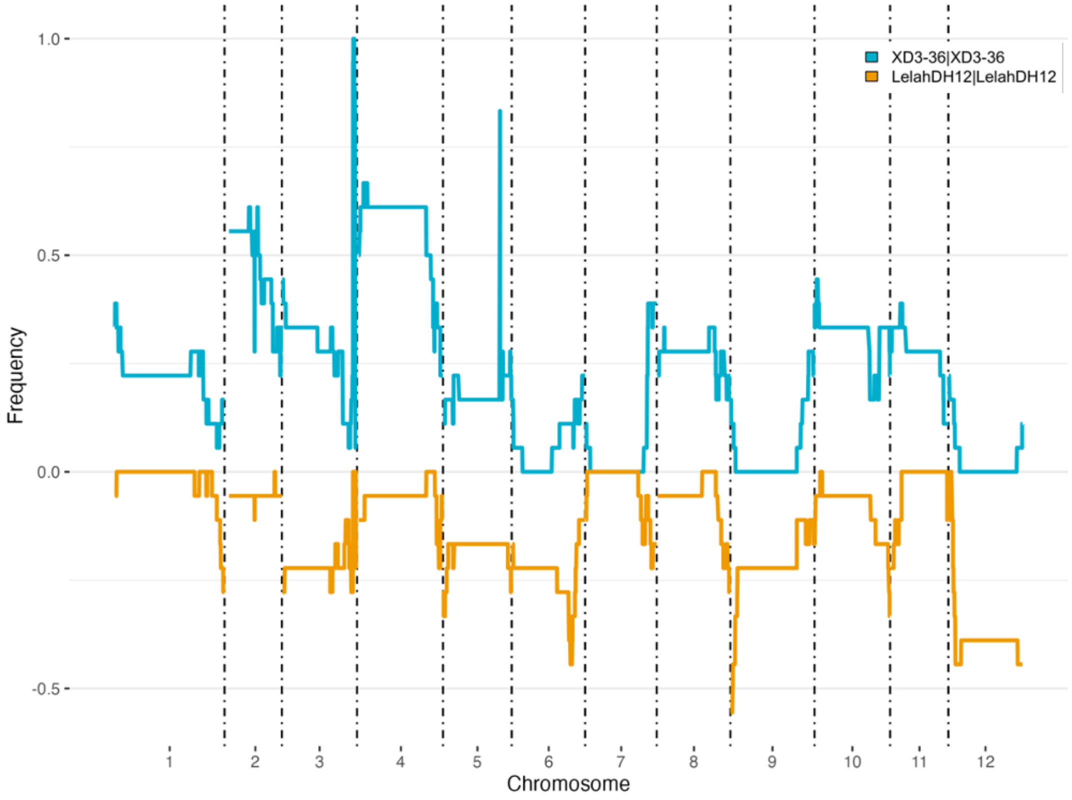


**Figure S3.** W2x001-22 $F_2$ haplotype frequency plot across 12 chromosomes for genotypes: XD3-36|XD3-36 (blue) and Lelah-DH12|Lelah-DH12 (orange).

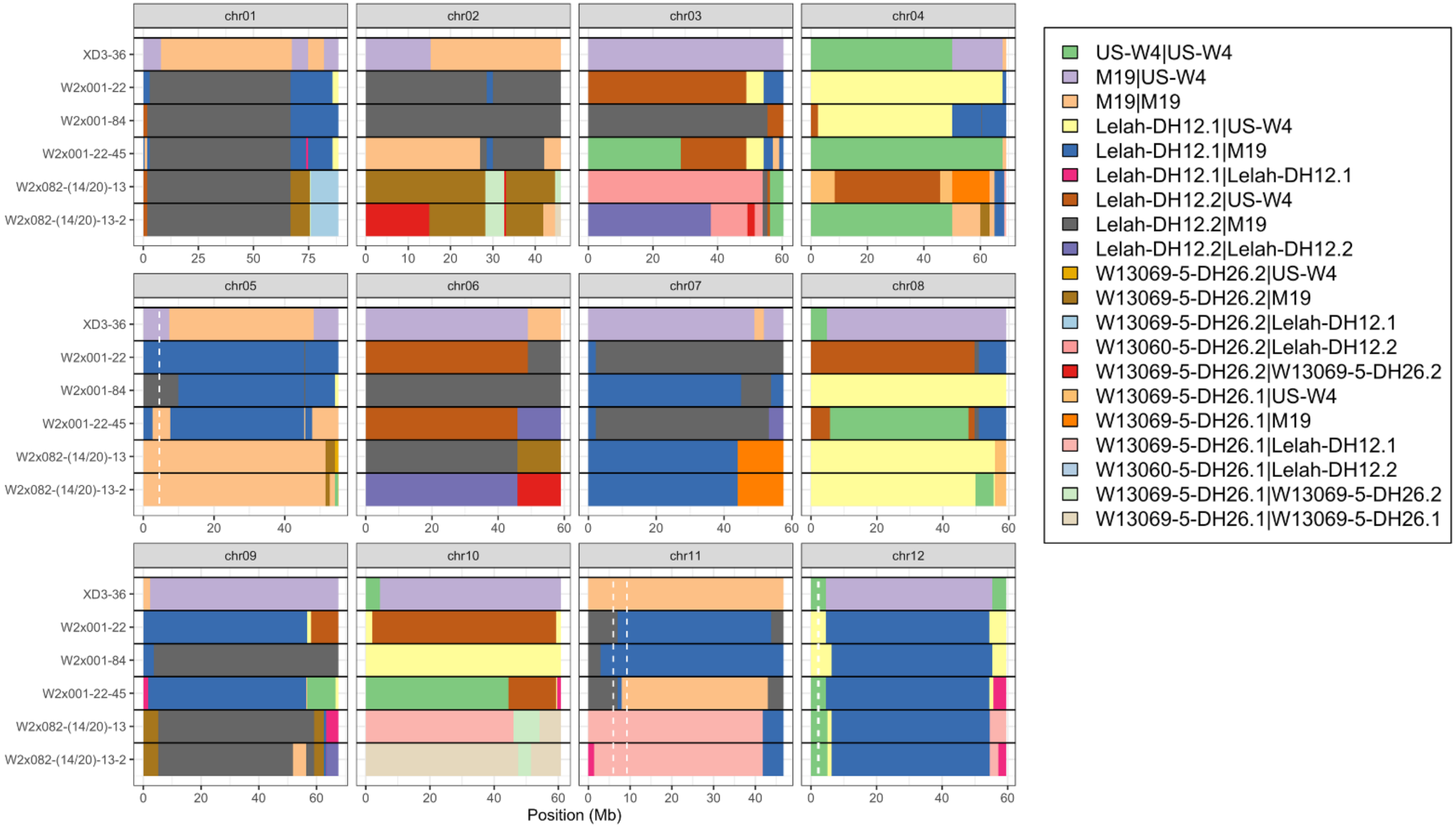


**Figure S4.** Haplotype reconstruction of key genotypes. White dashed lines represent the location of *CDF1* on chromosome 5, the fertility QTL on chromosome 11, and *Sli* on chromosome 12.

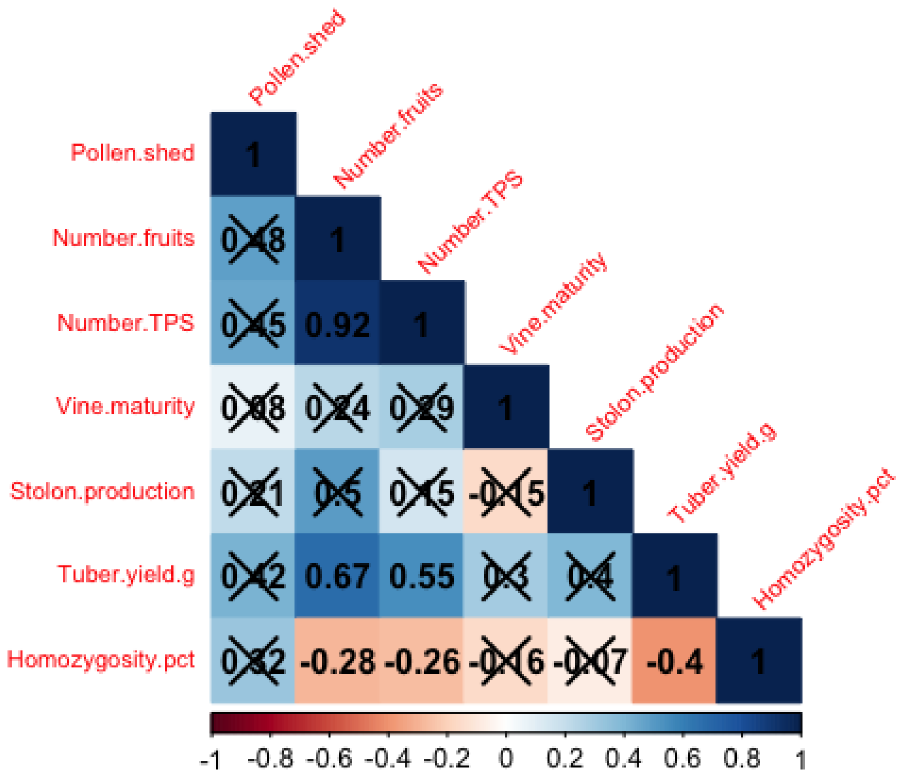


**Figure S5.** Correlation of traits for the W2x082-(14/20)-13-$F_3$ population. The correlation coefficients are calculated using Pearson's R for continuous-continuous cases, correlation ratio for categorical-continuous cases, and Cramer's V for categorical-categorical cases. "X" denotes a non-significant correlation at $\alpha$=0.05.

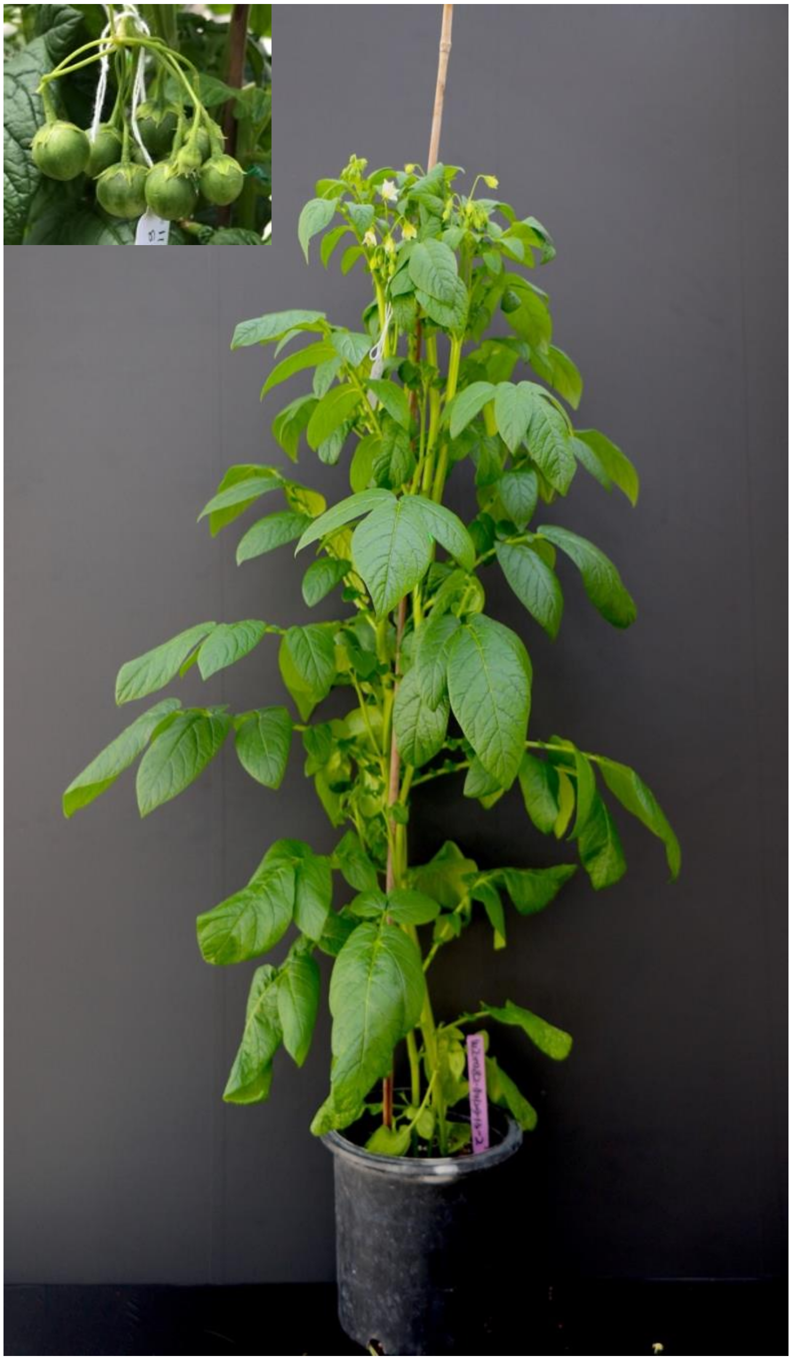

**Figure S6.** $BC_2F_3$ individual W2x082-(14/20)-13-2.
